# FLOE1 maintains cellular viscosity in rehydrating Arabidopsis embryos

**DOI:** 10.1101/2025.06.03.657678

**Authors:** Sterling Field, John F. Ramirez, Yanniv Dorone, Jack A. Cox, Thomas C. Boothby, Seung Y. Rhee

**Affiliations:** Plant Resilience Institute, Michigan State University, MI 48824, USA; Department of Biochemistry and Molecular Biology, Michigan State University, MI 48824, USA; Department of Plant Biology, Michigan State University, MI 48824, USA; Department of Plant, Soil, and Microbial Sciences, Michigan State University, MI 48824, USA; Department of Plant Biology, Carnegie Institution for Science, Stanford, CA 94305, USA; Department of Molecular Biology. University of Wyoming 1000 E. University Ave. Laramie, WY 82071

**Author notes:** Corresponding authors: Seung Y. Rhee;, Thomas C. Boothby.

**Keywords:** desiccation, seeds, biomolecular condensate, development, germination

## Abstract

Most plant embryos can survive for years in a dry state of less than 10% water(*1*). Rehydration during seed germination is critical but risky–too little environmental water can dehydrate and kill the developing embryo. Plants avoid this by germinating only when sufficient water is present. However, the mechanisms by which sufficient water is sensed and how it triggers germination remain unknown. FLOE1 suppresses seed germination under water limitation by undergoing reversible phase changes in response to water potential(*2*). To understand how FLOE1 affects germination, we compared water behavior in wild type and *floe1-1* null mutants. While both had similar amounts of water when dry, *floe1-1* seeds hydrated less than the wild type after imbibition. Additionally, bound water was less restricted in *floe1-1* embryos, suggesting FLOE1 promotes macromolecular water binding. By developing plant-compatible versions of Genetically Encoded Multimeric nanoparticles (GEMs)(*3*), we show that FLOE1 increases cytoplasmic viscosity during embryo rehydration. FLOE1 in yeast was sufficient to increase cytoplasmic viscosity. FLOE1 also increased cytoplasmic viscosity during hyperosmotic stress in yeast, and this ability was ablated by FLOE1 domain deletions. Our study identifies FLOE1 as a regulator of water content, dynamics, and cytoplasmic viscosity, linking molecular water control to cellular physical properties.

## Introduction

Cellular integrity, biochemistry and function depend on water availability as water is required for all metabolism. Cellular water influences membrane structure, protein folding, cellular crowding, and molecular interactions(*4–7*). As such, cells have evolved myriad means to modulate water content and behavior, ranging from the use of aquaporins to regulate water influx and efflux to the formation/dissolution of biomolecular condensates to tune water potential in response to environmental perturbations(*5*, *8*, *9*). A remarkable example of modulation of cellular activity linked to water content and potential is the ability of many plant embryos in seeds to reinitiate development only in the presence of sufficient water. Seeds of most plants can survive long-term in a dry state and water availability is critical for emergence from this dry, quiescent state and subsequent development(*10*). Developing embryos lose desiccation tolerance quickly upon rehydration, so too little water when the embryo of a seed reinitiates development may result in drying and death(*11*). To avoid this, many plants have evolved such that their embryos will only reinitiate development when sufficient water in the environment is sensed. However, historically, the mediator(s) involved in this “go or no go” developmental checkpoint were unknown.

Recently, it was shown that FLOE1, an intrinsically disordered protein abundant in developing plant embryos and seeds, acts as a regulator of germination in *Arabidopsis thaliana*(*2*). FLOE1 undergoes water-potential dependent changes in its physical phase, where under abundant water conditions, FLOE1 condenses into droplet-like puncta of 1-2 μm in diameter, whereas these condensates dissipate in water-scarce conditions, leading to a widespread distribution of FLOE1 throughout the cytoplasm of seeds upon drying(*2*). Targeted mutations that disrupt these phase changes, or lock FLOE1 in a particular phase, have corresponding effects on germination(*2*). Overall, FLOE1 governs germination of seeds via somehow sensing water potential and preventing premature reinitiation of germination in low water environments(*2*). Water behavior and dynamics are tuned by cellular components and their assembly/disassembly(*12*). Furthermore, water behavior affects cytoplasmic rheological properties(*12*, *13*). However, how plants leverage this tunability and whether this is a molecular mechanism underlying FLOE1’s ability to prevent precocious germination in low water conditions remains unknown.

Here, we aimed to uncover the molecular and cellular function of FLOE1 in preventing premature germination in low water environments. In rehydrating seeds, FLOE1 increases water content while simultaneously reducing water mobility. FLOE1 is also necessary for maintaining cytoplasmic viscosity during seed rehydration. We further assayed if FLOE1 was sufficient to alter cytoplasmic viscosity by expressing FLOE1 in a heterologous yeast system. FLOE1 was sufficient to increase cytoplasmic viscosity in yeast under hyperosmotic conditions. We then tested if FLOE1 domains, critical for phase separation, impact FLOE1’s modulation of cellular viscosity. Deleting the FLOE1 QPS domain increased viscosity in hyperosmotic conditions, while deleting the DS domain decreased cytoplasmic viscosity under these conditions. Together, these data lead to a proposed model in which FLOE1 regulates germination by facilitating water uptake, while keeping this water in a bound, immobilized state, such that it does not contribute to increasing water potential. Simultaneously, FLOE1 maintains high cytoplasmic viscosity, reducing the ability of embryonic constituents to vigorously diffuse and thus further restricts germination. To our knowledge, this work represents the first insights into how an organic molecule (e.g. sugar, lipid, protein, etc.) controls water dynamics to influence cytoplasmic material properties in the context of plant development. Beyond highlighting FLOE1’s significance in regulating biophysical properties during seed development and germination, our study offers prospects for novel bioengineering strategies for improving seed germination traits.

## Results

### FLOE1 increases water uptake and decreases molecular water mobility during rehydration

FLOE1 prevents precocious germination of seeds in low water environments(*2*), but how FLOE1 regulates germination at the molecular level is not yet understood. To address this, we quantified the total amount of water in *floe1-1* mutant seeds compared to wild type Col-0 seeds that were dry or imbibed using thermogravimetric analysis (TGA) (Fig 1A). TGA did not identify any difference in water content between *floe1-1* and Col-0 in the dry state (Fig. 1A). However, after 24 hr of imbibition, TGA analysis showed significantly increased water content in Col-0 relative to *floe1-1* seeds (Fig. 1A).

**Figure 1:**
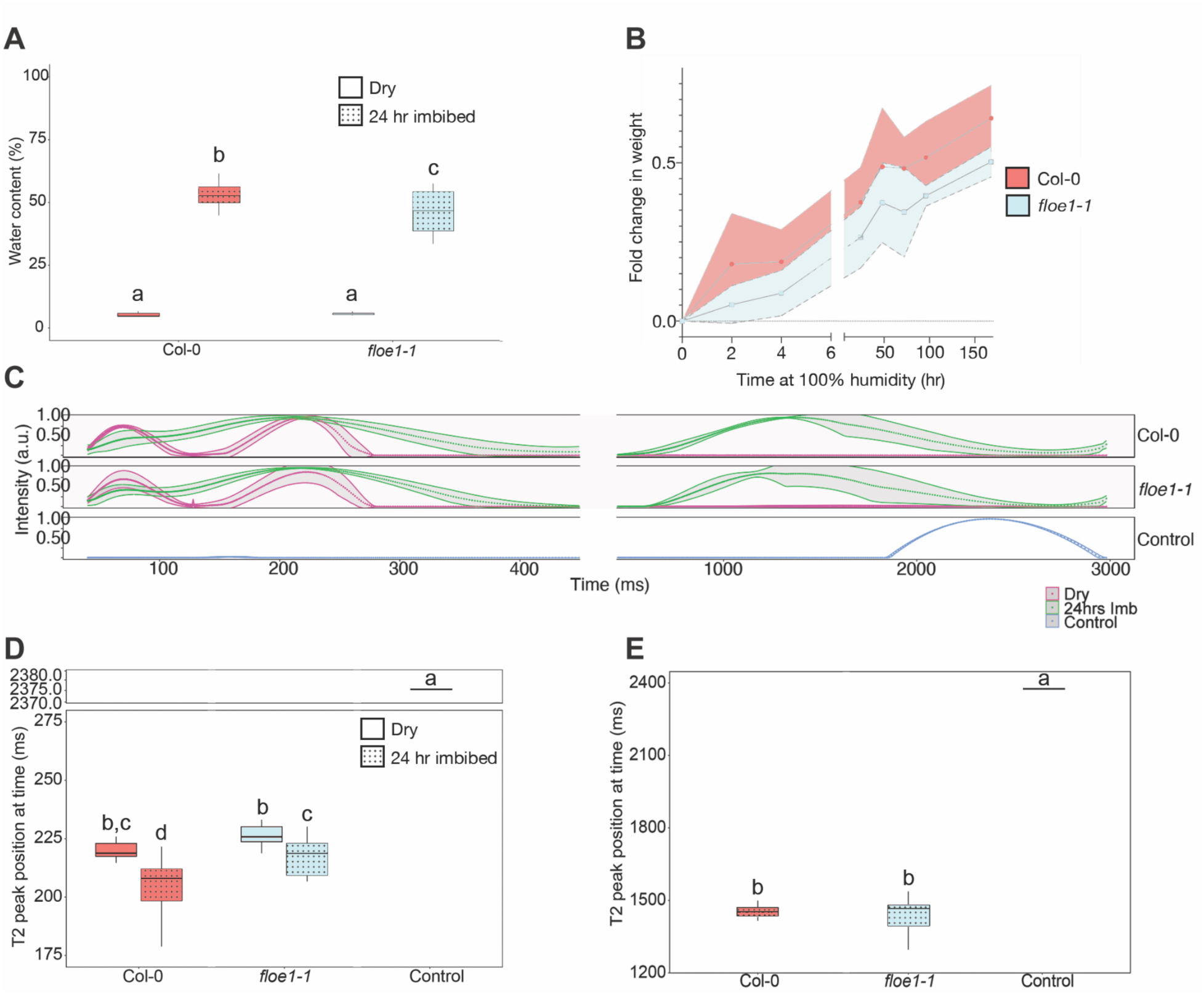
FLOE1 increases water content while simultaneously restricting water mobility in rehydrated seeds. (**A**) Water content in dry Col-0 and *floe1-1* seeds as well as after 24 hr of imbibition. (**B**) Water content in Col-0 and *floe1-1* seeds hydrated via humidification at 100% relative humidity for 150 hrs. Solid line denotes the average, while shaded areas denote the standard deviation. (**C**) Time-Domain NMR T2 values for pure water (blue) as well as Col-0 and *floe1-1* seeds in the dry (purple) and rehydrated (green) state. (**D**) T2 midpoint values for bound water (∼200 ms peaks) for Col-0 and *floe1-1* seeds in the dry and rehydrated state. Control is pure water. (**E**) T2 midpoint values for more mobile water (∼1,500 ms peaks) for Col-0 and *floe1-1* seeds after 24 hours of imbibition. Note these peaks are absent from dry samples (**C**) and thus not shown. Control is pure water. All statistics indicated are from two-way ANOVA with Tukey’s post hoc test.

TGA is a destructive assay used for measuring total water content at a single time point, but not the rate of water uptake. Therefore, we measured water uptake in *floe1-1* and Col-0 seeds by comparing the relative change in mass of seeds incubated in air saturated with water (100% relative humidity; RH) over the course of 150 hours (Fig. 1B). Similar to TGA analysis, water uptake at each time point was higher in Col-0 relative to *floe1-1* seeds (Fig 1B). Together, these data demonstrate that while FLOE1 does not influence the amount of water in dry seeds, it does increase the water content of seeds during rehydration.

One might intuitively assume that more water in wild type versus *floe1-1* seeds should mean that wild type seeds would germinate more quickly, However this is contrary to what was previously observed, where *floe1-1* seeds germinated more readily than Col-0 under low water potential(*2*). However, water does not exist solely in a fluid phase within a cell. Water binding to cellular constituents can lower the water potential of a system, thus limiting biological activity(*5*). With this in mind, we next asked if FLOE1 alters water dynamics.

We used Time-Domain NMR (TD-NMR) to assess if water in *floe1-1* and Col-0 seeds behaves as a bulk liquid, and if not, to what degree bound water is modulated by the presence of FLOE1. TD-NMR allows for the alignment of hydrogen nuclei spins and a subsequent measurement of the spin-spin relaxation time (T2)(*14*). For pure bulk liquid water, T2 is typically ∼2,300 milliseconds (ms), while non-bulk, less mobile water (e.g., water molecules bound to proteins) has lower T2 values(*15*, *16*).

Consistent with past literature, in our experiments, pure water had an average T2 of 2,378 ms (Fig. 1C-E). For dry Col-0 seeds, we identified a consistent T2 peak at ∼220 ms (Fig. 1C), which is in line with past TD-NMR work in seeds that identified “less mobile water” with this approximate T2(*17*). In dry *floe1-1* seeds, T2 for immobile water was ∼227 ms, which was significantly higher than dry Col-0 seeds (Fig. 1D, T-test, p≤0.001), demonstrating that the presence of FLOE1 is associated with less mobile water in dry seeds. We also observed a peak below 100 ms in both samples, which previous work attributes to oils in seeds(*17*). No higher peaks were identified in either of the dry samples, indicating, as expected, a lack of bulk fluid water (Fig. 1C).

To assess water dynamics upon rehydration, we imbibed Col-0 and *floe1-1* seeds for 24 hrs and then assessed T2 via TD-NMR (Fig. 1C-E). The difference in bound water mobility observed in dry seeds persisted after 24 hrs of imbibition, with Col-0 seeds having a significantly lower T2 (less mobile water) than *floe1-1* seeds (T-test, p≤0.005, Fig. 1C&D). As expected, imbibition caused the appearance of an additional peak at ∼1,450 ms, which we attribute to the accumulation of more mobile water within both genotypes (Fig. 1C&E). These more mobile water fractions were not significantly different between Col-0 and *floe1-1* seeds (T-test, p≥0.05, Fig. 1E). Combined, TD-NMR data indicate that in the dry state, as well as upon rehydration, bound water is less mobile in wild type seeds as compared to seeds lacking FLOE1.

Taken together, TGA and TD-NMR data indicate that FLOE1 increases total water uptake upon rehydration while simultaneously binding water and reducing its availability for biological processes(*18*). This behavior potentially represents an elegant means of accumulating water that will be needed for germination within the embryo while preventing precocious germination through sequestration of water.

### FLOE1 is necessary for decreasing cytoplasmic viscosity in germinating embryo

Sequestration of cytoplasmic water mediated by FLOE1 could increase cytoplasmic viscosity during rehydration, thus serving to slow the diffusion of cytoplasmic components and reducing rates of germination. With this hypothesis in mind, we wondered if FLOE1 might be a modulator of cytoplasmic viscosity.

To test this, we first employed classic rheological measurements using lysates obtained from 24 hr imbibed Col-0 and *floe1-1* seeds. We measured viscosity of both lysates under a range of shear stresses (Fig. 2A). The results showed that as expected at low shear, the viscosity of both samples was unaffected, followed by a region where increasing shear led to decreases in viscosity, and finally a region where additional stress no longer decreases viscosity. This pattern is modeled well by the Carreau-Yasuda model(*19–21*) for both our Col-0 and *floe1-1* lysates (R^2^ = 0.98 and 0.99, respectively; Fig. 2A). Fitting our data to this model allowed us to compare several key parameters. First we assessed the ‘zero shear viscosity’ (𝛈_0_) of both samples, which reports on the viscosity of each sample in the absence of any shear stress(*20–22*). Col-0’s zero shear viscosity was greater than that of *floe1-1* by more than three-fold (Fig. 2A&B), indicating that the presence of FLOE1 increases viscosity in rehydrating seeds. Next we quantified ‘onset shear rate,’ the amount of shear stress one must apply to a sample before its viscosity starts decreasing(*21*, *22*). Col-0 had an onset shear rate that was ∼30% higher than *floe1-1* (Fig. 2A&C), though this difference was not statistically significant. We also examined ‘endset shear rate,’ the total amount of shear stress needed for a sample to reach its minimum viscosity(*20*, *21*). Consistent with the zero shear viscosity, Col-0 seed lysates tolerated ∼3X as much shear stress before reaching their minimal viscosity compared to *floe1-1* seed lysates (Fig. 2A&D). Interestingly, the minimal viscosity (𝛈_∞_) obtained for each system was not significantly different, despite much more stress needing to be applied to Col-0 samples to reach that minimum (Fig. 2A&E). Taken together, these data indicate that the presence of FLOE1 increases the viscosity of rehydrating seed lysates.

**Figure 2:**
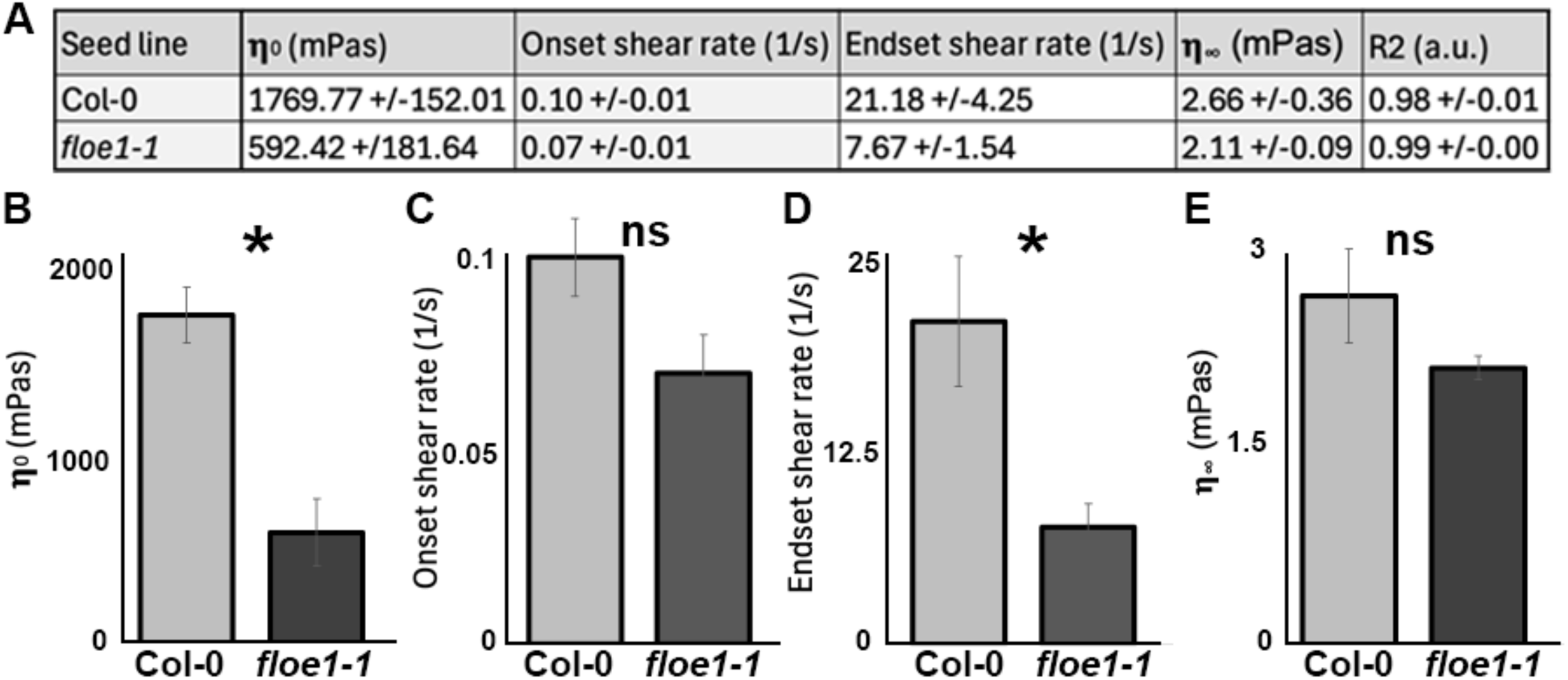
FLOE1 is necessary to maintain viscosity in seed lysates. (**A**) Summary table of key viscosity parameters: zero shear viscosity (𝛈_0_) (**B**), onset shear rate (**C**), endset shear rate (**D**), minimal viscosity (𝛈_∞_) **(E**), and the R2 value for the data’s fit with the Carreau-Yasuda model(*19–21*). (**B-E**) T-test, ns = not significant, * = p value of <0.05.

To determine how FLOE1 influences viscosity in the intact, live embryo, we next developed a method to assay changes in cellular viscosity in living plant cells. We opted for a live-cell imaging approach to measure cytoplasmic viscosity using Genetically Encoded Multimeric nanoparticles (GEMs)(*3*). GEMs are stable, 40 nm homomultimeric fluorescent particles previously used to track changes in cellular fluidic movements in yeast and mammalian cells(*3*). GEMs have not previously been optimized for use in plants, nor used previously to study changes in cytoplasmic viscosity during cellular desiccation and rehydration in any organism. Tracking the movement of GEM particles allows for calculation of diffusion rates, and serves as a way to measure viscosity at the subcellular level in living cells(*3*) (Fig. 3A).

**Figure 3:**
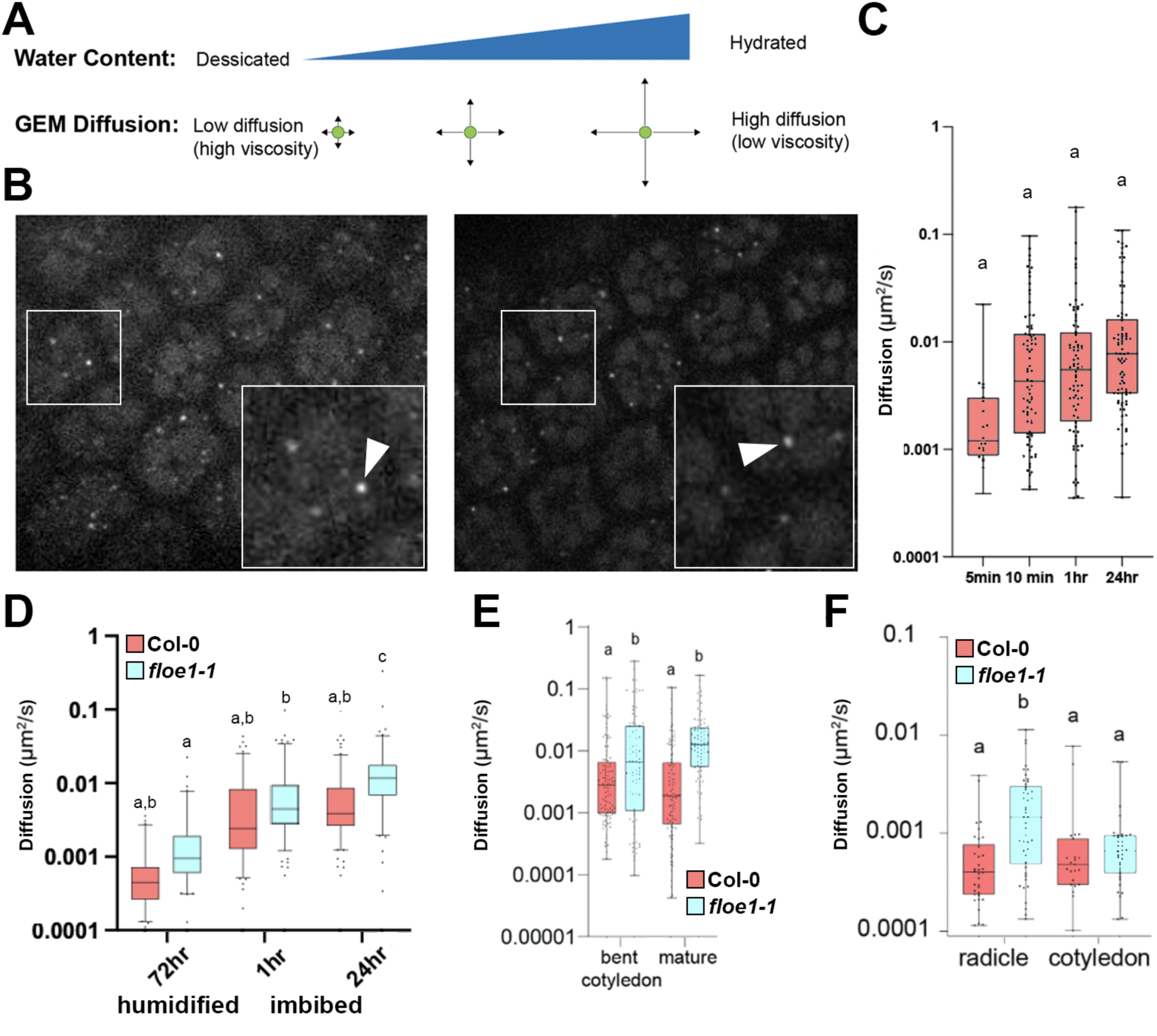
FLOE1 is necessary for maintaining cytoplasmic viscosity in imbibed seeds. (**A**) Predicted effect of hydration on GEM diffusion as a proxy for cytoplasmic viscosity. (**B**) Representative confocal Z-planes from the radicles of Col-0 (left) and *floe1-1* (right) embryos expressing GEMs (arrows). Larger boxed regions are enlarged inserts of smaller boxed regions. (**C**) Quantification of GEM diffusion in Col-0 embryos at different time points post imbibition. (**D**) Quantification of GEM diffusion in Col-0 (red) and *floe1-1* (blue) embryos after 72 hr of humidification (100% RH) as well as 1 and 24 hr of full imbibition. (**E**) Quantification of GEM diffusion in Col-0 (red) and *floe1-1* (blue) embryos at the bent cotyledon and mature stages of embryogenesis. Both the radicle and cotyledon were imaged. (**F**) Quantification of GEM diffusion in Col-0 (red) and *floe1-1* (blue) in the radicle and cotyledon of 24 hr imbibed embryos. (**C-F**) Statistical analysis used two-way ANOVA. Data is composed of at least 3 independent experiments, each independent experiment analyzing 3-9 cells.

We developed and characterized plant-optimized GEMs for expression in *Arabidopsis*. We used the published protein sequence for the 40 nm *Pyrococcus furiosus* coat protein codon optimized for *Arabidopsis* expression, driven by the constitutive *35S* promoter (GEMs/pGWB641) in stable *Arabidopsis* Col-0 and *floe1-1* lines (Fig. 3B). To assay GEMs in seeds, we first characterized GEM diffusion in imbibing Col-0 seeds (Fig. 3B; Movie S1). GEMs in Col-0 readily formed upon addition of water and allowed us to measure diffusion rate starting after 5 minutes of imbibition (Fig. 3C). In Col-0, cytoplasmic viscosity did not significantly increase over the course of 24 hrs, though the median diffusion rate did change from ∼0.001 to ∼0.009 um^2^/s (Fig. 3C).

Using the GEMs, we asked if the cytoplasm of *floe1-1* seed cells was less viscous (i.e. more diffusion occurring) than Col-0 during germination (Fig. 3D). Seed rehydration and germination is a dynamic process where cells experience close to no water (∼5-15% water) and end with being fully rehydrated (∼90% water)(*23*, *24*). We started by assessing cytoplasmic viscosity in *floe1-1* and Col-0 seeds that had been humidified at 100% RH for 72 hr, which leads to slow diffusion of water into the seeds. While the mean diffusional rate was greater in *floe1-1* seeds relative to Col-0, this difference was not significant (Two-way ANOVA, p=ns, Fig. 3D). Next, we assayed the diffusion rates of GEMs in *floe1-1* seeds after 1 and 24 hr of imbibition, water diffuses into the seeds much faster, and compared them to Col-0 (Fig. 3D). In 1 hr imbibed seeds, Col-0 and *floe1-1* had similar cytoplasmic GEM diffusion rates (Fig. 3D). However, GEM diffusion rates in *floe1-1* and Col-0 seeds behaved differently when comparing 72 hr at 100% RH to 1 hr imbibition (Fig. 3D). For Col-0, GEM diffusional rates after 72 hr at 100% RH and 1 hr imbibition were indistinguishable from each other. However, *floe1-1* seeds imbibed for 1 hr had significantly higher GEM diffusion rates than *floe1-1* seeds exposed to 100% RH for 72 hr (Two-way ANOVA, p≤0.05, Fig. 3D). Furthermore, cytoplasmic diffusion increased significantly in *floe1-1* at 24 hr versus 1 hr (Two-way ANOVA, p≤0.0001), whereas diffusion in Col-0 seeds did not change significantly between 1 and 24 hr (Two-way ANOVA, p=ns, Fig. 3D). Finally, at 24 hr of imbibition diffusion rates in *floe1-1* seeds were significantly higher than in Col-0 seeds (Two-way ANOVA, p≤0.0001, Fig. 3D). Using diffusion of GEMs as a proxy for cytoplasmic viscosity (more diffusion corresponding to less viscosity), we conclude from these experiments that the presence of FLOE1 attenuates the decrease in cytoplasmic viscosity upon rehydration of the seed - that is seeds with FLOE1 maintain viscosity longer after imbibition relative to seeds lacking FLOE1.

### FLOE1 is necessary for increasing cytoplasmic viscosity in dehydrating embryo during seed maturation

Opposite of seed rehydration and germination, seeds undergo dehydration during seed maturation, a process that must be controlled to enable desiccation tolerance and seed longevity(*25*). It is yet unknown what controls this process. Therefore, we were curious whether FLOE1 influences changes in cytoplasmic viscosity during seed maturation. To address this, we assessed GEM diffusion rates in *floe1-1* and Col-0 plants at the early-maturation (bent cotyledon) and mid-maturation (seed filling) stages of seed development(*26*) (Fig. S1). Both the early- and mid-maturation stages of *floe1-1* embryos displayed greater diffusional rates compared to Col-0 embryos (Fig. 3E). These data indicate that the presence of FLOE1 mediates the reduction of cytoplasmic viscosity during seed maturation.

An additional advantage of using GEMs to assess molecular diffusion and cytoplasmic viscosity is the ability to obtain these parameters at cellular and tissue resolutions. In 24 hr imbibed *floe1-1* seeds, most of the decrease in cytoplasmic viscosity was observed in cells in the embryonic root (radicle), while cytoplasmic viscosity of cotyledon cells remained similar to Col-0 (Fig. 3F). The viscosity of the radicle and cotyledon in Col-0 seeds was indistinguishable (Fig. 3F). These observations suggest that FLOE1 works in a tissue specific fashion, specifically reducing cytoplasmic viscosity in the radicle of *Arabidopsis* seeds after 24 hr imbibition.

Taken together, these data demonstrate that FLOE1 is necessary for attenuating decreased cytoplasmic viscosity in imbibed seeds and increases cytoplasmic viscosity in dehydrating seeds during seed maturation. Furthermore, it appears that FLOE1 acts in a tissue specific fashion, reducing viscosity in the radicle, but not the cotyledons.

### FLOE1 is sufficient to increase cytoplasmic viscosity under hyperosmotic conditions

After observing that FLOE1 is necessary for attenuating viscosity decrease in the imbibed embryo, we next asked if FLOE1 was sufficient to increase viscosity. To test this, we used *Saccharomyces cerevisiae* (baker’s yeast), which does not harbor the *FLOE1* gene. We first engineered a galactose induced GEM-mEmerald construct for expression in yeast cells. To test this construct, we assayed GEM diffusion in wild type (W303a) yeast under standard conditions (Fig. 4A). The diffusion of GEMs in the cytoplasm of wild type yeast was ∼0.1 um^2^/s (Fig. 4A), corroborating previous estimates(*3*). To further confirm that GEMs were behaving as expected in our system, we tested how cytoplasmic diffusion is impacted when yeast cells are treated with a brief (10 minute) incubation with low-osmotic pressure NaCl solutions, as hyperosmotic shock is known to increase viscosity(*27*) (Fig. 4A). As expected, wild type yeast exposed to increasing levels of hyperosmotic conditions displayed increasing viscosity (Fig. 4A).

**Figure 4:**
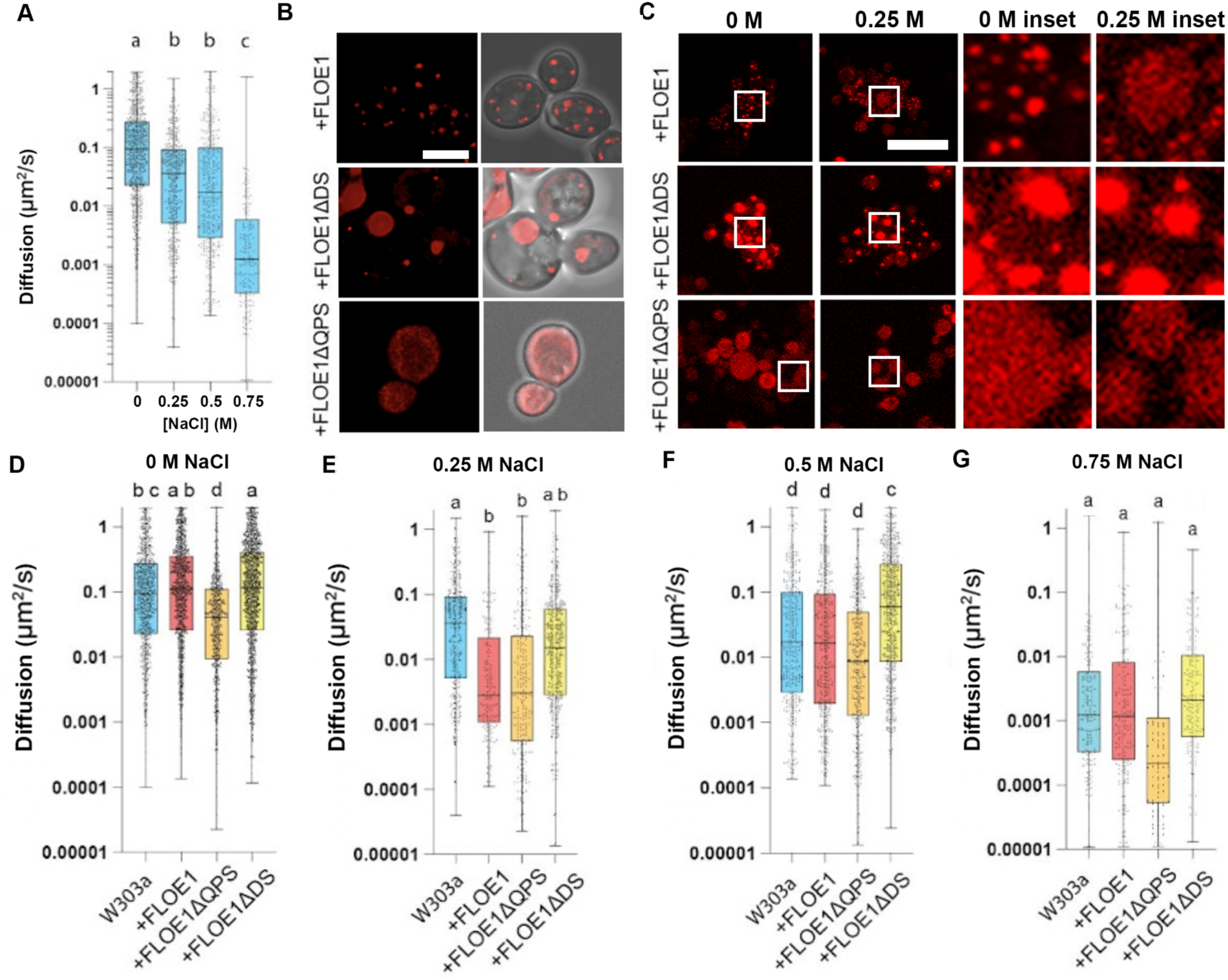
FLOE1 is sufficient to increase cytoplasmic viscosity during hyperosmotic stress. (**A**) Quantification of GEM diffusion in wild type W303a yeast with increasing concentrations of NaCl. (**B**) Confocal imaging showing the localization FLOE1-mCherry, FLOE1ΔDS-mCherry, and FLOE1ΔQPS-mCherry in yeast under normal (left) and hyperosmotic (right) conditions. (**C**) Confocal imaging showing the localization FLOE1-mCherry, FLOE1ΔDS-mCherry, and FLOE1ΔQPS-mCherry in yeast under normal and hyperosmotic stress conditions (250 mM NaCl). White boxes indicate inset areas. (**D**) Quantification of GEM diffusion in different yeast strains under normal osmotic conditions, (**E**) with 0.25 M NaCl, (**F**) 0.5 M NaCl, and (**G**) 0.75 M NaCl. (**A, D-G**) One-way ANOVA.

After observing that GEMs corroborate previous estimates of cytoplasmic viscosity and respond to changes in water potential in our system, we next assessed the effect of FLOE1 on GEM diffusion. We generated a transgenic yeast line in our W303a-GEM background expressing FLOE1. Consistent with its behavior in *Arabidopsis* seeds(*2*), FLOE1 condensed into puncta in hydrated yeast (Fig. 4B&C) and these puncta dispersed upon hyperosmotic shock (Fig. 4C). In addition, we generated two additional yeast lines containing domain deletions of FLOE1. We previously identified two domains of FLOE1 important for FLOE1 function in regulating germination in low water conditions, suggesting they may be critical for FLOE1 function(*2*). The FLOE1 QPS domain is an intrinsically disordered region, which when deleted, results in a FLOE1 null phenotype(*2*). Similar to results from *Arabidopsis*(*2*), FLOE1ΔQPS did not form puncta when expressed in yeast under normal or hyperosmotic conditions (Fig. 4B&C). The second domain, DS, is predicted to have single alpha helical domain flanked by disordered regions and deletion of this domain results in larger, gel-like FLOE1 body that no longer disperses under high water potential conditions and leads to precocious germination under low water potential conditions(*2*). Consistent with its behavior in *Arabidopsis*(*2*), FLOE1ΔDS formed larger condensates under normal and hyperosmotic conditions in yeast(Fig. 4B&C).

To assess whether FLOE1 is sufficient to reduce cytoplasmic viscosity, we compared GEM diffusion in wild type and FLOE1-expressing yeast cells under standard osmotic conditions (Fig. 4D). We found no significant difference in diffusion between wild type and FLOE1-expressing yeast under these conditions (Fig. 4D). We next asked whether QPS and DS deletions would affect diffusion under standard conditions. For FLOE1ΔDS-expressing cells, which exhibit constitutively condensed FLOE1, we observed no significant differences in diffusion relative to wild type or FLOE1-expressing yeast (Fig. 4D). However, FLOE1ΔQPS-expressing yeast cells, which exhibit constitutively dispersed FLOE1, reminiscent of wild type FLOE1 during water deficit, we observed reduced diffusion relative to wild type, FLOE1, and FLOE1ΔDS yeast cells under standard conditions (Fig. 4D). These results demonstrate that FLOE1 does not affect cytoplasmic viscosity in yeast when water potential is high, but might increase viscosity during water deficit when FLOE1 exists in a dispersed, decondensed state.

To test if FLOE1 is sufficient to increase cytoplasmic viscosity during water deficit, we assessed GEM diffusion in our yeast cells exposed to increasing levels of hyperosmotic stress (Fig. 4E-G). As expected, we observed a general trend where hyperosmotic shock led to increased viscosity in all yeast strains regardless of the presence of FLOE1 (Fig. 4D-G). For example, diffusion decreased by two orders of magnitude from ∼0.1-0.015 um^2^/s without NaCl to ∼0.001-0.00015 um^2^/s with 0.75 M NaCl (Fig. 4D&G). Under standard conditions, FLOE1-expressing cells did not alter diffusion rates compared to wild type W303a cells (Fig. 4D). However, under mild hyperosmotic shock (0.25 M NaCl), FLOE1-expressing cells decreased their GEM diffusion rates by more than one order of magnitude compared to W303a cells (Fig. 4E). Interestingly, at greater hyperosmotic stress this difference in diffusion between wild type and FLOE1 expressing yeast became non-significant (Fig. 4F&G). This convergence on similar diffusional rates between FLOE1-expressing and wild type cells was driven by a steady decrease in diffusion within wild type cells while diffusion in FLOE1-expressing cells remained invariant and low at all levels of hyperosmotic stress (Fig. 4E-G).

Since FLOE1 reduces water mobility (Fig. 1C&D), we were concerned that this might be inducing a hyperosmotic response, which is yeast is mediated by the high osmolarity glycerol (HOG) pathway, which in turn might be leading to reduced cytoplasmic viscosity(*28*). To this end, we quantified the diffusion of GEMs in yeast lacking HOG1, a gene encoding a mitogen-activated protein kinase involved in osmoregulation and required for the HOG pathway(*29*). Knockout of the HOG1 gene in the W303a background did not lead to a significant change in diffusion compared to wild type W303a (Fig. S2). Similarly, knockout of HOG1 in the FLOE1 expressing strain did not lead to significant changes in diffusion compared to yeast expressing FLOE1 alone (Fig. S2). Our HOG1 knockout experiments demonstrate that activation of the HOG pathway via heterologous expression of FLOE1 is not a major contributor to the cytoplasmic viscosity reduction observed in FLOE1 expressing lines.

Collectively, these observations demonstrate that FLOE1 is sufficient to enhance cytoplasmic viscosity when water becomes limiting. Increased cytoplasmic viscosity by further dehydration ultimately surpasses the degree to which FLOE1 increases viscosity, supporting the notion that in *Arabidopsis* seeds, the presence of FLOE1 serves to maintain high cytoplasmic viscosity during seed germination.

Consistent with our observations that loss of the FLOE1 QPS domain constitutively recapitulates the behavior of wild type FLOE1 during hyperosmotic stress in *Arabidopsis*, we observed that diffusion in FLOE1 and FLOE1ΔQPS yeast strains were not significantly different under hyperosmotic conditions (Fig. 4D-F). Conversely, we observed that FLOE1ΔDS yeast cells did not reduce diffusion more than wild type W303a cells under any hyperosmotic conditions (Fig. 4D-E). To assess whether phase separation of any protein will induce changes in viscosity, or whether the changes in viscosity we observe are due to the phase of FLOE1 specifically, we measure GEMs diffusion in yeast expressing the *A. thaliana* RHAMNOSE1 enzyme (RHM1). RHM1 is involved in the synthesis of uridine diphosphate (UDP)-glucose and is known to undergo phase separation(*30*). Expression of RHM1 in yeast did not significantly increase or decrease diffusion of GEMs (Fig. S3), indicating that while phase separation of FLOE1 tunes cytoplasmic viscosity, phase separation in general does not. These results demonstrate that the ability of FLOE1 to increase viscosity under hyperosmotic stress is linked to the ability of the protein to phase separate.

## Discussion

The ability of many seeds to desiccate and remain dormant until favorable conditions are present for germination has contributed to their successful colonization of every continent and nearly every terrestrial biome. However, emergence from this dry, quiescent state is precarious. If a plant initiates embryogenesis and development in an environment without sufficient water, it risks drying out and dying. How seeds sense water has been an enigma for centuries, until recently, when it was shown that FLOE1 could govern germination of *Arabidopsis* seeds through changing its biophysical state in response to water potential change(*2*). However, how FLOE1 functions to reduce germination rates under water deficit was not known.

Here, we show that FLOE1 functions as a modulator of water content, water dynamics, and cytoplasmic viscosity. We show that FLOE1 simultaneously increases water content while reducing the mobility of water within imbibed seeds. This ‘immobilization’ of water is concomitant with increased cytoplasmic viscosity that is FLOE1-dependent. Furthermore, FLOE1 is sufficient to induce increased viscosity in heterologous systems under conditions of water deficit and this ability is linked to the capacity of FLOE1 to undergo a phase change from a condensed (associated with reduced cytoplasmic viscosity) to dispersed (associated with increased cytoplasmic viscosity) state.

Combining the results of this study with Watson and colleagues’ theory on water dynamics and molecular condensation(*5*), we propose a mechanism for how FLOE1 changes cytoplasmic viscosity to elicit the physiological changes in the Arabidopsis embryo to germinate. Our previous work highlighted an intriguing dynamic behavior of FLOE1, which is hydration dependent(*2*). In the dry state, FLOE1 is dispersed throughout the cytoplasm, while rehydration leads to the molecular condensation of the protein(*2*). This behavior, combined with insights from a previous study on water dynamics and molecular condensation(*5*), provides an elegant potential means for FLOE1, in its dispersed state, to bind to and sequester greater amounts of water without increasing water potential or jumpstarting embryonic development. Upon sufficient hydration, FLOE1 condenses into droplets, with this condensation liberating previously bound water as FLOE1-water interactions are replaced by FLOE1-FLOE1 interactions. The liberated water is then available to promote germination. Thus, through a delicate interplay between water content and dynamics, cytoplasmic viscosity, and protein phase behavior, FLOE1 serves to regulate germination of the embryo through the accumulation of inactive water early during rehydration and subsequent release of active water. This sequestration/release cycle of water is mirrored by changes to the material and biophysical properties of the cytoplasm, which FLOE1 helps to keep in a super-viscous state. We propose that these mechanisms coalesce to provide a safeguard against precocious development in environments that are prone to redrying.

## Methods

### Arabidopsis

#### Growth conditions

*Arabidopsis* plants were grown under long day conditions (16 hours light, 8hrs dark, at 21°C, at ∼50% humidity, with a light intensity of 150 µmol/m^2^).

#### Molecular Cloning

The *Arabidopsis* GEMs construct used the 40 nm GEMs amino acid sequence derived from *Pyrococcus furiosus* coat protein (PDB ID: 2E0Z), cloned into pGWB641 (GenScript). GEMs/pGWB641 was transformed into *Arabidopsis thaliana* using (*31*).

#### Microscopy

GEMs imaging. To assay GEM diffusion rates, seed cells were visualized using a Zeiss Axiovert + Yokogawa CSU-W1 T1 spinning disk confocal microscope. GEMs were visualized with a 63x glycerol immersion objective, using a 488 laser (laser intensity set to 45%) and detecting fluorescence between 510 nm - 530 nm. GEM movement was imaged by capturing 100 images, taken at a rate of 15 frames per second. Images were saved as .sld files. Image analysis for GEMs is further explained at the end of the methods section.

#### Preparing tissues for water weight and GEM assays

##### Seed humidity assays

Mature dry seeds for seed weight and GEM analysis were incubated in 100% humidity conditions. To make 100% humidity conditions, water soaked paper towels were placed in a plastic tupperware box, and allowed to acclimate for 24 hrs. 100% humidity was confirmed using a digital hygrometer. To incubate seeds, seeds were first aliquoted into eppendorf tubes, and quickly placed into the plastic tupperware box (box was open less than 15 seconds). After incubating for 24 hr, seeds were then analyzed.

##### Imbibe assays

Mature dry seeds for seed weight and GEM analysis were imbibed in water by first aliquoting seeds into eppendorf tubes, and then adding excess distilled water to the seeds.

##### Dissecting embryos for GEMs assays

Developing *Arabidopsis* plants were grown as stated above. Maturing siliques were cut off *Arabidopsis* plants, and under a dissecting microscope the siliques were cut lengthwise, exposing the developing embryos. The developing embryos were then placed on glass slides and sorted visually into early-maturation (bent cotyledon) and mid-maturation (seed filling) groups (see Supplemental Figure 1), followed by imaging, as described above.

##### Seed weight assays

For seed weight assays, 100 *Arabidopsis* seeds were placed into a previously weighted PCR tube, and the total weight was measured, yielding the weight of 100 dry *Arabidopsis* seeds. Seeds were then incubated at 100% humidity. At timepoints listed in the graphs, the humidity incubated seeds were directly weighted.

#### Seed water content data (TGA)

For measurement on TGA approximately 0.5 mL of samples were placed into 1.5 mL centrifuge tubes and labeled for imbibed conditions. 1.0 mL of distilled water was added to tubes. Tubes were inverted and vortexed repeatedly to ensure water surrounding all seeds. Tubes were then kept on benchtop room at temperature for 24 hours. After 24 hours samples were removed from tubes with scupula and spread thinly across Kimwipes to absorb excess water prior to running on TGA. For dry and imbibed condition seeds, a scupula was then used to collect seeds and place on pre-tared TGA crucibles.

Samples were run on a TA TGA5500 instrument in 100 μL platinum crucibles (TA 952018.906). Crucibles were tared prior to each run and prior to sample loading. Crucibles were loaded with between 5 mg and 10 mg of sample mixture. Each sample was heated from 30 °C to 250° C at a 10 °C per minute ramp. All TGA data and thermograms are available in Supplemental File 2 and Supplemental File 3.

Determination of water loss was conducted using TA’s Trios software (TA Instruments TRIOS version #5.0.0.44608). Thermograms were used to calculate starting masses of samples and the mass of samples at the plateau that occurs after ∼100 °C but before the thermal denaturation at ∼200 °C. The Trios software “Smart Analysis” tool was used to identify the inflection point between these two mass loss events.

#### TD-NMR

For Time Domain NMR measurement approximately 1 mL of seeds were placed into 10 mm flat bottom borosilicate Time Domain NMR Tubes (Wilmad-LabGlass WG-4001-7). Dry condition sample tubes were immediately capped after the addition of seeds. Imbibed condition samples then had 8 mL of distilled water added to each TD-NMR tube and were capped. Tubes were inverted and vortexed repeatedly to ensure water surrounding all seeds. Tubes were then kept on a benchtop at room temperature for 24 hours. After 24 hours, 7 mL of water was removed from each tube. Afterwards, an additional 1 mL of water and seeds was removed from each tube leaving 1 mL of water and seeds in each tube. Imbibed condition tubes were then recapped.

T_2_ relaxation measurements were performed using a Bruker mq20 minispec low-field nuclear magnetic resonance spectrophotometer, with a resonance frequency of 19.65 MHz. Samples were kept at 25°C during measurements through the use of a chiller (F12-MA, Julabo USA Inc., Allentown, PA) circulating a constant-temperature coolant. T_2_ free induction decays were measured using a Carr-Purcell-Meiboom-Grill (CPMG) pulse sequence with 8000 echoes for dry samples and 6000 for imbibed samples, and an echo time of 1000 μs. Pulse separation of 0.25 ms was used for dry samples and 1.5 ms for imbibed samples, recycle delay of 3 ms, and 8 scans were used for all samples. The gain was determined for each sample individually via software (The minispec Software: V3.00 Rev.10) automated tune-gain. Conversion of raw TD-NMR data to NMR spectra was performed with Open Source 1D/2D Inverse Laplace Program (OSILAP, Takahiro Ohkubo; https://amorphous.tf.chiba-u.jp/memo.files/osilap/osilap.html).

#### Rheometry

Rheometry measurements were performed on a TA HR 20 instrument with Relative Humidity Accessory. Samples were run in a parallel plate geometry on stainless steel, 40 mm disposable plates (TA 527400.907). The Relative Humidity Chamber was preset with the desired set points of 23°C and 50%RH. The parallel plate geometry had a running gap of 1000.0µm and a trim gap of 1050.0µm. Measurements were performed via logarithmic sweep of shear rate from 0.01 1/s to 500.0 1/s with 5 points per decade collected. Seed lysates were measured by placing lysates onto clean bottom surface plates and then excess seed lysate was trimmed from the outside of the plates after achieving the trim gap. The Relative Humidity Accessory chamber was then sealed and allowed to reach the preset temperature and humidity. After measurement the plates were wiped clean of debris and lysate after every run with Kimwipes and ethanol.

Seed lysates were prepared using 15mL glass douncers were used to homogenize seeds for collection of seed lysates prior to running rheometry experiments. Prior to homogenization seeds were allowed to imbibe for 24 hours. Excess water was removed from seeds via patting dry with Kimwipes. To produce adequate seed lysate volume 25.2 mg of seeds and 0.5 mL of ultrapure water were added into end of the douncer. 40 consecutive cycles consisting of applying pressure, turning handle 180°, and pulling a vacuum before resetting to original position and pressure. After 40 consecutive cycles of homogenization, the douncer was removed from douncer body and an additional 1 mL of ultrapure water added to assist collection of seed lysate. A pipette was then used to transfer the seed lysate to a labeled 1.5 mL microcentrifuge tube. The microcentrifuge tubes were then centrifuged at maximum speed for 1 minute to pellet seed debris. Seed lysate supernatant was then run in parallel plate geometry on stainless steel, 40 mm disposable plates.

Analysis of logarithmic sweep of shear rate was performed by TA’s TRIOS software (TA Instruments TRIOS version #5.6.0.87). The TRIOS software “Best fit flow (viscosity vs. rate)” tool was used to determine best model fit and perform analysis of data utilizing the best fit model.

### Yeast

#### Molecular Cloning

GEMs for yeast work used 40 nm GEMs derived from *Pyrococcus furiosus* coat protein (PDB ID: 2E0Z) fused in frame with mEmerald (*Gal*:GEM-mEmerald/pESC-TRP; GenScript). For FLOE1 and the FLOE1 domain deletions, a yeast-codon optimized FLOE1 was synthesized and cloned into pESC-URA (GenScript), and the domain deletions were generated using site directed mutagenesis (GenScript) on the FLOE1/pESC-URA plasmid.

#### Growth conditions

*Saccharomyces cerevisiae* (strain W303a; ATCC 208352) cells were grown in Difco YPD Broth (#242820). To transform W303a cells with expression constructs, W303a cells were transformed with FLOE1/pESC-URA, FLOE1ΔDS/pESC-URA, FLOE1ΔQPS/pESC-URA, or GEMs/pESC-TRP following the Frozen EZ Yeast Transformation II Kit protocol (T2001, Zymo). Yeast was initially grown on SD-URA (FLOE1/pESC-URA, FLOE1ΔDS/pESC-URA, and FLOE1ΔQPS/pESC-URA), SD-TRP (GEMs/pESC-TRP), or SD-TRP/URA media with 2% Dextrose at 30°C. SD media was supplemented with 1 x yeast nitrogen base without amino acids and ammonium sulfate (Difco #233520) and 2 % dextrose, or by replacing dextrose with 2 % galactose (final concentration; Sigma, CAS-No 59-23-4) to induce protein expression. For experiments, yeast cultures were pelleted (3,000 x g for 3 minutes), and resuspended in either 0 M, 0.25 M, 0.5 M, or 0.75 M NaCl. For control experiments, a similar approach was used but using either BY4772 or BY4772-ΔHOG1.

#### Microscopy

##### Imaging FLOE1 bodies

FLOE1-mCherry and FLOE1 domain deletion mutants in yeast were visualized using a Leica SP8 scanning laser confocal microscope (Leica, Wetzlar, Germany). Proteins tagged with mCherry were excited with a white light laser at 561 nm and fluorescence was detected in the range of 574 nm - 629 nm by a hybrid detector.

##### Imaging GEMs in yeast

To assay GEM diffusion rates, cells were visualized using a Zeiss Axiovert + Yokogawa CSU-W1 T1 spinning disk confocal microscope, or a Leica Thunder Fluorescent microscope. For visualizing GEMs using the Zeiss microscope, GEMs were visualized using a 488 nm laser (laser intensity set to 25%), and fluorescence was detected at 510 nm - 530 nm and a 100x glycerol immersion objective. Using the Thunder Fluorescent Microscope, GEMs were visualized using a 485 nm laser (20% laser intensity) and detected with a GFP filter and K8 camera, using a 100x oil objective. GEM movement was imaged by capturing 100 images, taken at a rate of 20 frames per second.

#### GEMs analysis

To generate diffusion coefficients from GEMs images, images of GEMs were first imported and opened in FIJI (version 1.0). Images were then analyzed using the MOSAICsuite plugin(*32*), and each stack of 100 images was analyzed using the Mosaic built-in Particle Tracker function. To first detect particles in images, we used a threshold of 0.3 percent of absolute image intensity to detect particles. Next, we tracked particles that had a link range of 15 (i.e. only particles visible in 15 sequential frames were measured). Diffusion coefficient data from each individual GEM was then compiled in excel, and diffusion rates were graphed in Prism (version 10.3.0).

## Statistics

T-test, one-way or two-way ANOVA with Tukey’s post hoc test was used to compare treatments as indicated in figure legends. Graphs and statistical tests used Prism (GraphPad; version 10.3.0).

## Data presentation

Unless otherwise noted, all graphs present the full range of data points. For statistical significance, different letters represent a p-value ≤ 0.05.

## Acknowledgements

We would like to thank members of the Rhee and Boothby labs for helpful discussion on this project. We would like to thank Andrey Malkovskiy at the Carnegie Institution for Science Microscopy Core for assistance with microscopy. The ΔHOG1 yeast strain was a gift from Prof. Hugo Tapia (CSUCI). This work was done in part on the ancestral land of the Muwekma Ohlone Tribe which was and continues to be of great importance to the Ohlone people, and on the ancestral, traditional, and contemporary lands of the Anishinaabeg – Three Fires Confederacy of Ojibwe, Odawa, and Potawatomi peoples.

This work was funded in part by the U.S. National Science Foundation grants (IOS-2312181, IOS-2406533, IOS-1546838, MCB-1617020, and OISE-2434687 to SYR, DBI-2213983 to SYR and TCB) and U.S. Department of Energy, Office of Science, Office of Biological and Environmental Research, Genomic Science Program grants (DE-SC0018277, DE-SC0020366, DE-SC0023160, DE-SC0008769, and DE-SC0021286 to SYR). We thank the members of WALII for their insights and helpful discussions related to this work.

## Author contributions

Conceptualization: SF, SYR, JFR, TCB

Methodology: SF, SYR, JFR, TCB

Investigation: SF, SYR, JFR, TCB, JAC

Visualization: SF, JFR

Resources: YD

Funding acquisition: SYR, TCB

Project administration: SYR

Supervision: SF, SYR, TCB

Writing – original draft: SF, SYR, JFR, TCB

Writing – review & editing: SF, SYR, YD, JAC, JFR, TCB

## Competing interests

Authors declare no competing interests.

## Materials & Correspondence

Plant and yeast materials should be requested from Seung Rhee

## Supplementary information

**Supplementary movie 1:** GEMs expressed in Col-0. Mature and dry Col-0 seeds were imbibed in water, after 1 hr the embryo was dissected out of the seed and visualized by fluorescent microscopy. Video is of cells in the cotyledon.

### Supplementary Figure and Legends

**Supplementary Figure 1:**
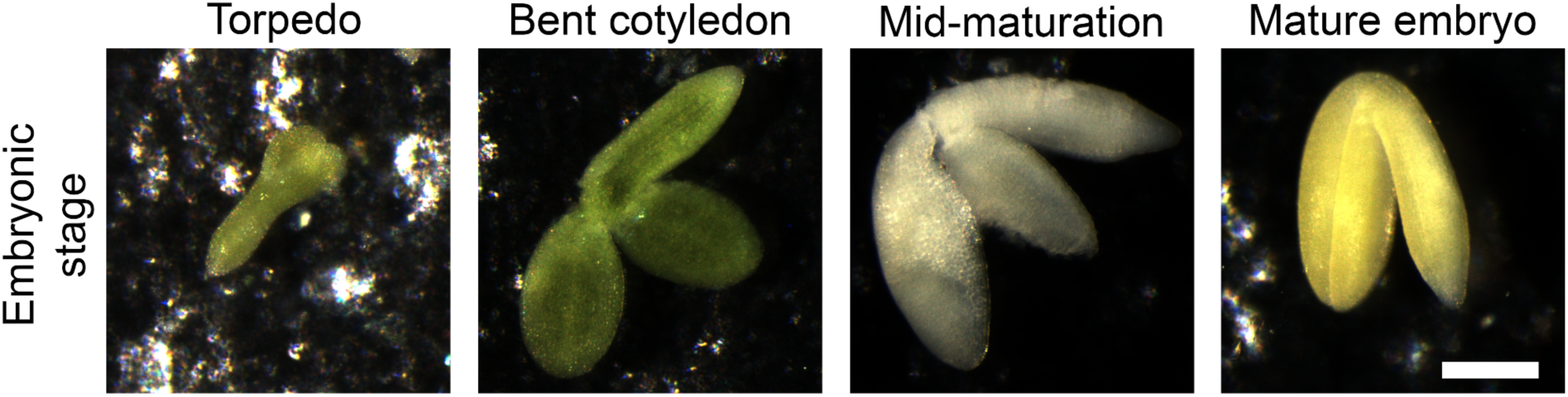
*A. thaliana* embryonic stages. GEMs diffusion was assessed in the bent cotyledon and mid-maturation stages of embryogenesis.

**Supplementary Figure 2:**
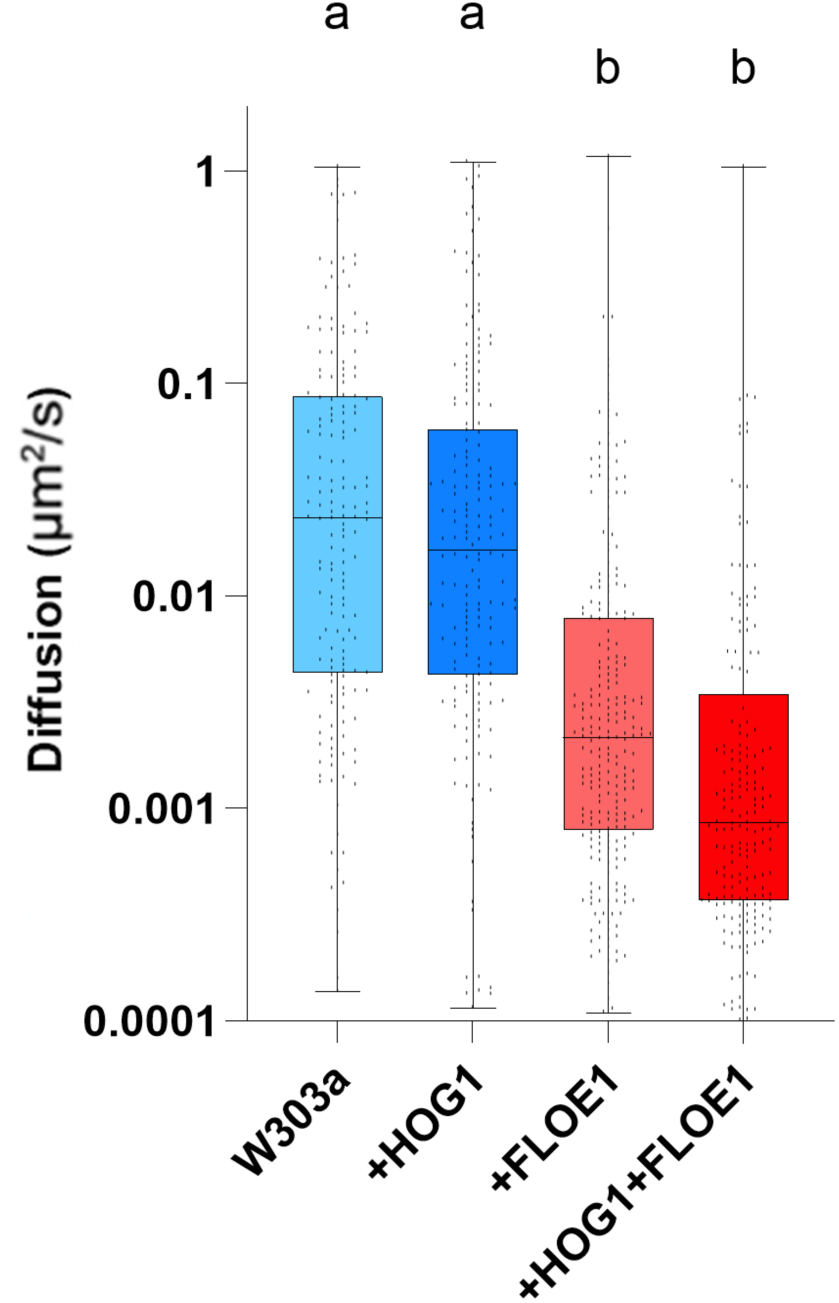
Reduced diffusion is not linked to activation of the high osmolarity glycerol (HOG) pathway. GEMs diffusion in yeast strains exposed to 250 mM NaCl. One-way ANOVA with Tukey’s post hoc test.

**Supplementary Figure 3:**
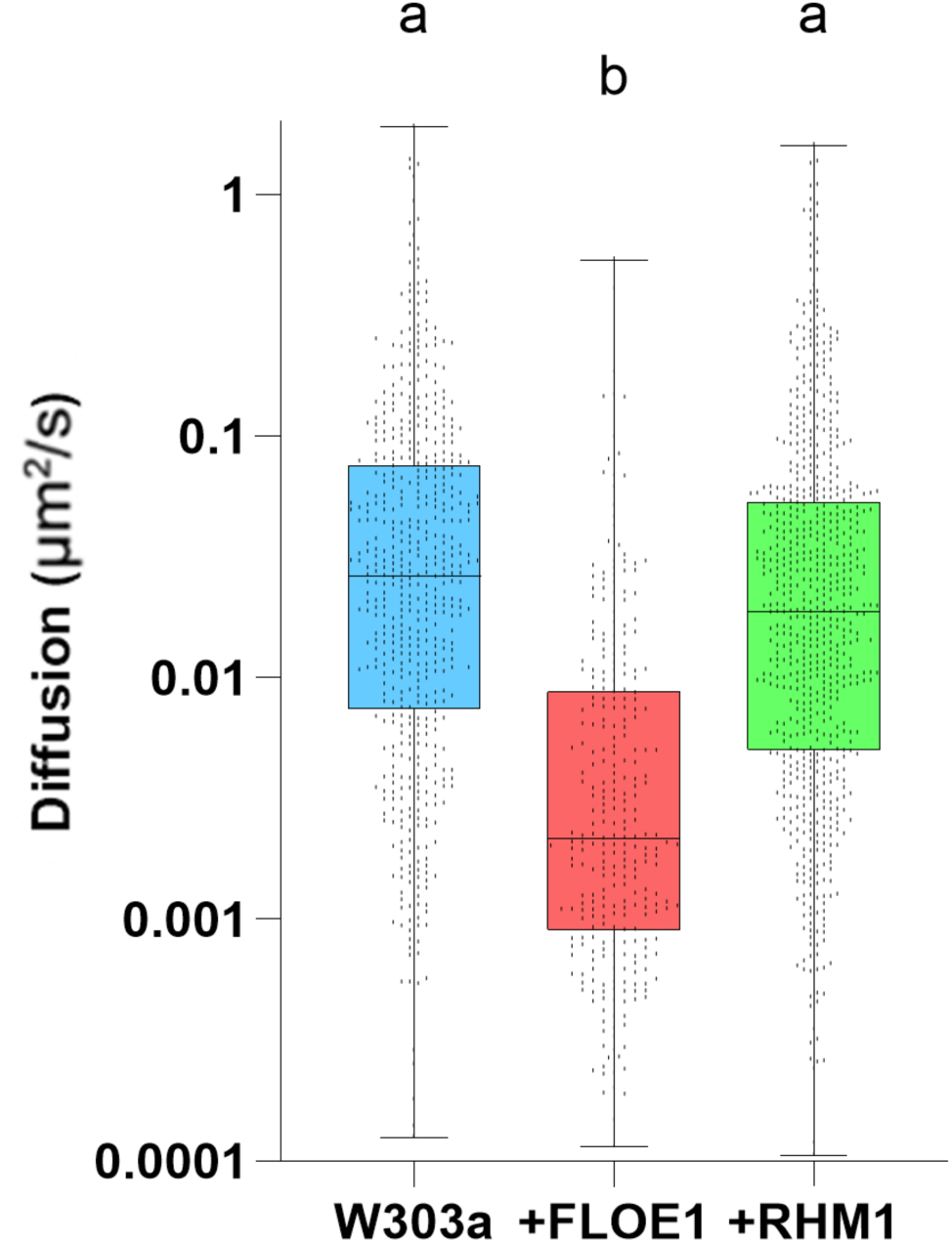
Heterologous expression of the phase separating protein RHM1 does not reduce diffusion in yeast. GEMs diffusion in yeast strains exposed to 250 mM NaCl. One-way ANOVA with Tukey’s post hoc test.

## Notes

### Competing Interest Statement

The authors have declared no competing interest.

